# Curvature-Encoded Photochemistry for Programmable Isolation and Fractionation of Extracellular and Plasma Membrane Vesicles

**DOI:** 10.64898/2025.12.30.697083

**Authors:** Jennifer Pichurin, Advaith Nair, Amelia Williams, Sikha Thoduvayil, Abhijith Radhakrishnan, Sudhanshu Gautam, Aditi Naskar, Pallavi Gopal, Themis Kyriakides, Sathish Ramakrishnan, Roshan W. Gunasekara

## Abstract

Lipid-based nanoparticles, including extracellular vesicles (EVs), cell membrane vesicles (CMVs), and biomimetic nanovesicles, are emerging as powerful carriers for therapy and diagnosis due to their ability to transport diverse payloads across biological barriers. EVs, in particular, show promise for liquid biopsy and systemic therapy because of their endogenous origin, biocompatibility, and intrinsic targeting capacity. Cell-derived plasma membrane nanovesicles complement EVs by preserving the source cell’s membrane architecture and receptor landscape, creating customizable scaffolds for biomimetic drug delivery. However, the full potential of these lipid nanovesicles has been challenging because current isolation workflows are labor-intensive, low-yield, and often lack purity, forming a critical bottleneck for clinical translation and standardized manufacturing. To overcome these challenges, we developed a programmable electrostatic and light-activated platform for isolating lipid vesicles directly from complex biological fluids in a rapid, high-purity, and scalable manner. Our approach employs photocleavable lipid nanoprobes (PLNs) that insert into vesicle membranes in a size-dependent fashion and, together with a cationic polymer, enable efficient capture on a solid support. A brief light exposure cleaves the probes to trigger on-demand release and fractionation of intact vesicles with high speed and precision. Using this platform, we demonstrate high-purity EV isolation with preserved membrane protein functionality, as well as single-step enrichment of right-side-out (RSO) cell membrane nanovesicles that maintain native membrane orientation for effective drug delivery. The capability to rapidly isolate and tailor vesicle subpopulations by size and orientation establishes a unified materials platform for precision nanomedicine and advanced therapeutics, paving the way toward clinical-grade vesicle production and next-generation liquid biopsy applications.

## 1. Introduction

Lipid-based nanovesicles have emerged as a versatile class of soft functional materials at the interface of synthetic nanotechnology and cell-derived therapeutics.^[1, 2^] This family encompasses extracellular vesicles (EVs), cell membrane vesicles (CMVs), and biomimetic liposomes, all built around a lipid bilayer that can host and protect molecular cargo.[3–5] Their high surface-to-volume ratio, modular membrane composition, and compatibility with chemical and biological surface engineering enable precise control over pharmacokinetics, biodistribution, and cellular targeting.[6, 7] In principle, such designable vesicle systems could integrate drug delivery, gene modulation, and minimally invasive diagnostics within a single platform of nanomaterials. Realizing this vision requires innovative isolation and fractionation strategies capable of delivering vesicle populations with high purity, defined size, and preserved membrane orientation at throughput scales compatible with clinical translation.

Within the broader class of lipid vesicles, EVs and CMVs are particularly attractive because they inherit the molecular signatures and targeting motifs of their source cells. EVs circulate throughout the system and are currently being investigated for purposes such as liquid biopsies, immune modulation, and targeted drug or gene delivery.[8–10] In parallel, CMVs derived from cell plasma membranes retain the antigenic and functional landscape of their parent cells, offering unique biomimetic advantages such as extended circulation, immune evasion, and homotypic targeting.[11–13] For example, RBC-derived vesicles leverage surface proteins such as CD47 to evade immune clearance, while tumor-derived vesicles exploit homotypic targeting to enhance specificity in cancer therapy.[14–16] Biomimetic liposomes further expand this toolkit by allowing complete control over lipid composition, size, charge, and release properties.[17] Despite this enormous potential, translating vesicle-based technologies to the clinic has been impeded by the lack of efficient, scalable purification methods.

Clinical deployment of EVs, CMVs, and related nanovesicles is fundamentally limited by current isolation and purification techniques, particularly at the scales required for therapy or manufacturing. EVs are typically harvested from complex biological fluids, such as cell culture media and blood plasma, which are rich in proteins, lipoproteins, and cellular debris. Likewise, standard liposome synthesis often produces heterogeneous vesicle sizes and leaves residual starting materials or byproducts. Removing these contaminants while preserving vesicle integrity and yield is non-trivial. Conventional workflows, including ultracentrifugation, density gradients, size-exclusion chromatography, and tangential-flow filtration, are laborious, time-intensive, and difficult to scale. They often expose vesicles to high shear or centrifugal forces and multiple handling steps, which can deform or damage the vesicles, change their cargo, and prevent precise control over size or membrane orientation. Crucially, existing methods offer little to no ability to distinguish right-side-out (RSO) vesicles with native membrane orientation from inside-out (ISO) vesicles. This lack of refinement in isolation techniques is a significant obstacle for applications requiring uniform, orientation-specific vesicle preparations, such as delivering membrane proteins that must face outward to interact with target cells or producing vesicles within a specific size range for uniform biodistribution.

Emerging methods aim to address different aspects of this challenge. Lipid-based probes that insert into vesicle membranes via hydrophobic anchors have been studied for their ability to selectively capture vesicles and remove non-vesicular contaminants.[18–22] However, most probe-based techniques still rely on affinity tags or chemical ligands for binding and release, which introduce additional steps and make scaling up more difficult. Adding DNA tags to lipid probes could enable controlled release via enzymatic digestion of PEG linkers and affinity handles.[18] However, this approach is too expensive and complex to implement efficiently at a large scale in manufacturing. To meet the need for large-scale, affordable, high-purity vesicle isolation, there is a demand for a platform that provides rapid capture, programmable release, and control over subpopulations in a simple, easily scalable format.

In this work, we present a next-generation, programmable, electrostatic, and light-activated purification system that enables rapid, high-purity, and scalable isolation of lipid vesicles while preserving their integrity and biological functionality. Our method employs photochemistry to release capture tags following vesicle isolation. It also pairs this with a cationic polymer to create a robust yet reversible electrostatic attraction. This allows for the selective capture of negatively charged vesicles, such as EVs and CMVs, from complex biological samples. Photocleavable lipid nanoprobes spontaneously insert into vesicle membranes in a size-dependent manner: smaller vesicles incorporate fewer probes, whereas larger vesicles contain more. A brief, adjustable light exposure then acts as a trigger to distinguish and release vesicle populations based on size. The combination of rapid PLL-mediated capture and highly efficient photocleavage allows the platform to operate on clinically relevant volumes within minutes while maintaining vesicle integrity. This system also enables enrichment of RSO-oriented vesicles, directly addressing the long-standing challenge of orientation-based separation. By addressing major challenges in vesicle purification, such as speed, purity, scalability, size-selective release, and orientation control, our platform provides a transformative solution for producing clinical-grade nanovesicles for diagnostics and therapy.

## 2. Results and Discussion

### 2.1. Development of a Programmable Electrostatic and Light-Activated (PELA) platform for rapid isolation of small extracellular vesicles

We developed a Programmable Electrostatic and Light-Activated (PELA) Platform to address the limitations of traditional EV isolation methods, thereby enhancing scalability and maintaining vesicle integrity. The system includes a photocleavable lipid nanoprobe (PLN) and positively charged poly-L-lysine (PLL) polymer. As depicted in Figure 1A, the PLN is an amphiphilic molecule with a hydrophobic lipid tail for membrane insertion, a UV-sensitive linker for light clevage, a PEG spacer to prevent non-specific binding, and a desthiobiotin affinity tag for anchoring to monomeric avidin beads. This multifunctional setup enables dual-mode vesicle isolation via light- or affinity-based capture. We used a 25:75 (PLL:PLN) ratio as optimal (Figure 1B) composition for efficient vesicle engagement through electrostatic attraction and hydrophobic membrane insertion without impeding subsequent photorelease. Notably, this coupling of PLL-mediated rapid capture with photocleavable release is central to the platform’s ability to operate efficiently in large-volume workflows using low-cost reagents and simple benchtop workflow.

**Figure 1.**
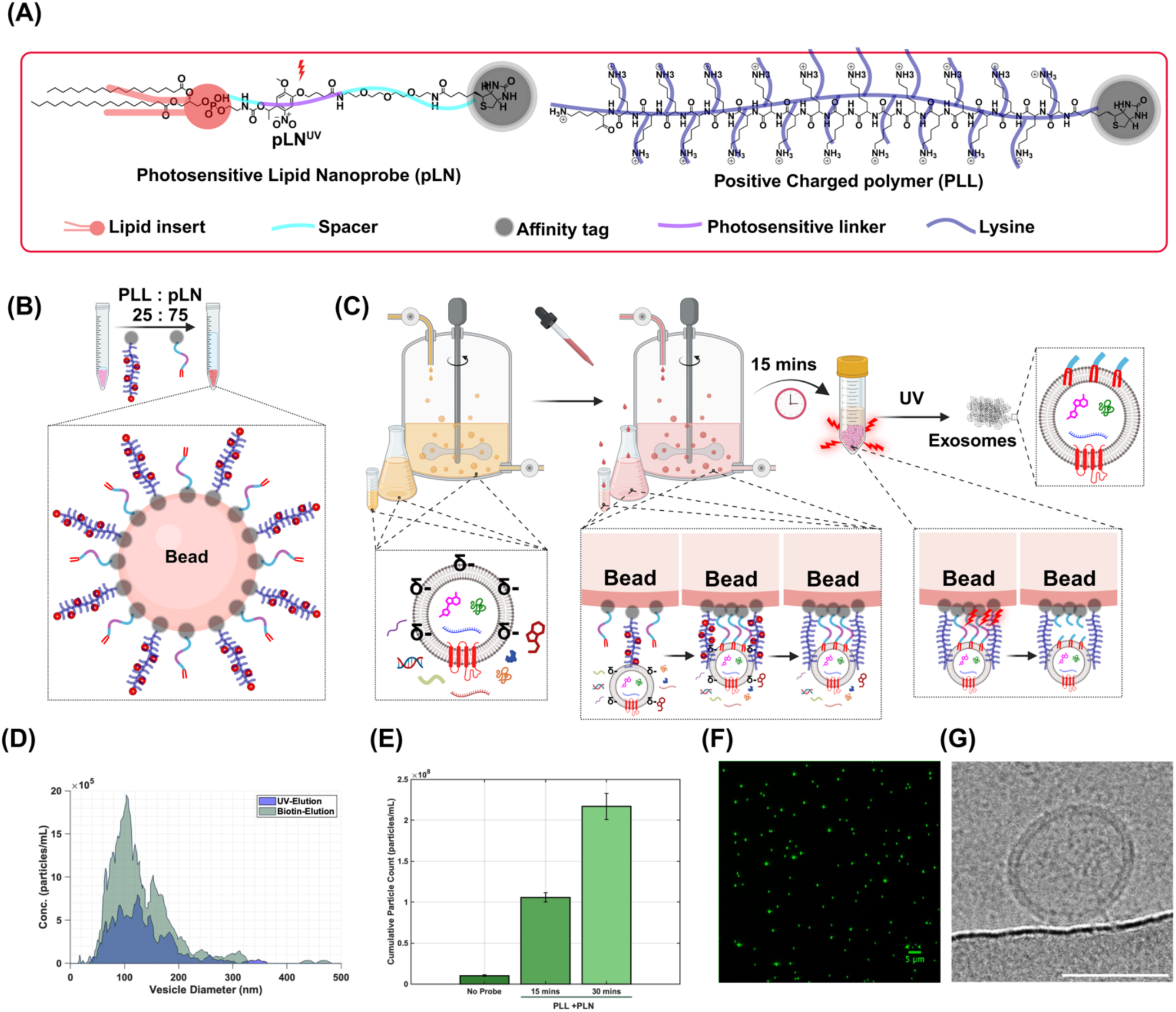
Programmable Electrostatic and Light- Activated (PELA) Platform for Rapid Isolation of Extracellular Vesicles (EVs) for Large-Scale Applications. (A) Schematic of the PELA components: a photocleavable lipid nanoprobe (PLN) and a positively charged polymer (PLL). PLN comprises a hydrophobic lipid insert for EV membrane integration, a UV-sensitive linker (red), a spacer, and an affinity tag (grey circle), enabling dual-mode isolation. PLL facilitates electrostatic interactions with negatively charged EV membranes. (B) Schematic of the PELA platform containing PLL and PLN at a 25:75 ratio. (C) Schematic of PELA-mediated EV isolation workflow. The PELA complex binds EVs via electrostatic and hydrophobic interactions. Subsequent UV irradiation cleaves the PLN linker, releasing intact EVs while removing non-vesicular contaminants. (D) Nanoparticle tracking analysis (NTA) demonstrates comparable size distributions of EVs isolated via UV- or biotin-mediated elution. (E) Quantitative analysis of EV capture efficiency demonstrates significantly higher EV capture within 15 and 30 minutes with the PELA platform than with controls lacking PLN, highlighting rapid and efficient isolation. (F) Fluorescence microscopy images of EVs labeled with MemGlow-488 membrane dye showing successful isolation after 15 mins of PELA platform incubation. (G) Representative cryo-transmission electron microscopy (cryo-TEM) image of an isolated EV, revealing intact, spherical morphology (Scale bar: 50 nm).

Using this design, we established a streamlined EV isolation workflow (Figure 1C). Briefly, the PELA formulation is introduced into conditioned cell culture medium, enabling PLNs to embed into vesicle membranes while PLL promotes rapid vesicle association. The probe-functionalized vesicles are captured on the solid support through the PLN affinity handle, followed by extensive washing to remove non-vesicular contaminants that do not carry PLNs. A short pulse of UV light then cleaves the PLN linker, disconnecting the vesicles from the immobilized probe and releasing intact EVs into solution. The overall workflow is completed in less than 15 minutes, eliminating the need for ultracentrifugation or lengthy column elutions. By avoiding harsh physical forces and extensive handling, this workflow preserves the native structure and cargo of vesicles, thereby enabling a scalable transition from small laboratory samples to large-volume clinical or industrial samples.

We next benchmarked vesicle output by comparing UV-triggered photorelease with competitive affinity elution using D-biotin. NTA revealed highly similar size distributions between these two modes of recovery, with a predominant peak at ∼100 nm (Figure 1D), indicating that the light-based release does not measurably deform or damage EVs. We then quantified capture efficiency from 50 mL of conditioned medium. PELA achieved significantly higher EV capture within 15 minutes relative to controls lacking PLNs (Figure 1E), demonstrating that the combined electrostatic attraction (PLL) and membrane insertion (PLN) enables rapid and efficient vesicle scavenging from large volumes.The capture yield increases for a longer period, underscoring that the combination of PLNs and PLL rapidly and effectively scavenges vesicles from solution. These results highlight the speed and efficiency of our approach for high-throughput vesicle isolation.

We also confirmed that the isolated vesicles remain structurally intact using microscopy. Fluorescence imaging of membrane-stained EVs using Memglow-488 dye revealed bright, punctate signals after just 15 minutes of PLD-mediated capture, indicating successful concentration and recovery of vesicles (Figure 1F). Consistent with these results, cryo-TEM imaging revealed EVs with intact, spherical morphology and clearly defined lipid bilayers (Figure 1G). These observations illustrate that our gentle, light-based release preserves the native vesicle architecture, making it suitable for downstream applications that require functional vesicles.

### 2.2 PLN Probes preserve the vesicle morphology and membrane protein functionality

We next asked whether PLN-mediated capture and UV-triggered release compromise vesicle structure or membrane protein activity. To evaluate both parameters in a controlled and quantitative manner, we generated biomimetic extracellular vesicles (bEVs) that combine native EV components with defined synthetic lipids (Figure 2A) using two fabrication routes: (i) extrusion and (ii) direct mediated reconstitution/dialysis with defined membrane protein content for functional assays. The extrusion-based approach enables incorporation of synthetic lipids into EV-derived membranes while maintaining EV-like morphology and providing tight control over vesicle size distributions.

**Figure 2.**
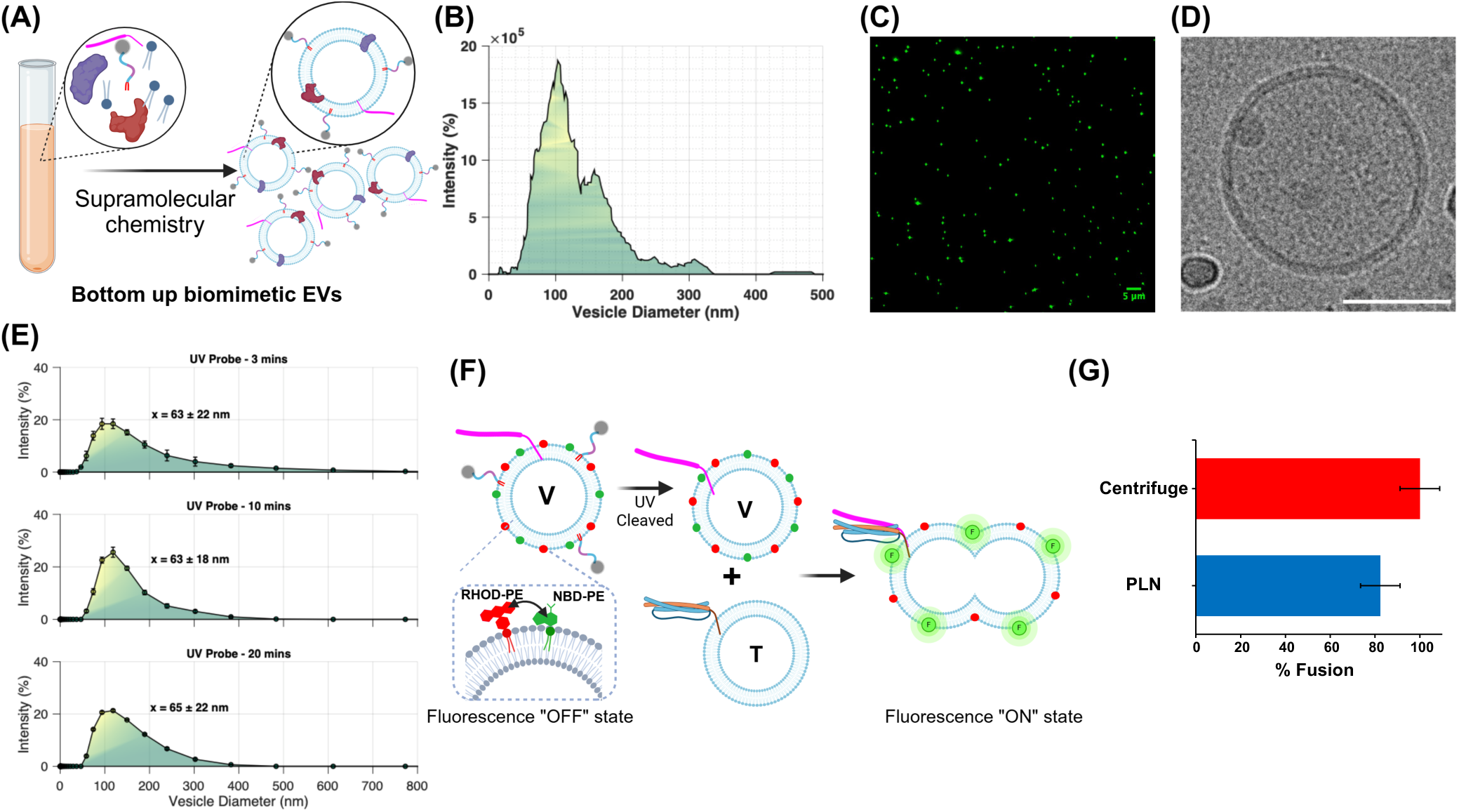
Rapid Isolation and Functional Characterization of Biomimetic EVs Fabricated via a Bottom-Up Approach. (A) Schematic illustrating the bottom-up assembly of biomimetic EVs incorporating SNARE proteins from defined molecular components. (B) Representative NTA profile of isolated biomimetic EVs. (C) Fluorescence microscopy image of isolated vesicles, confirming successful isolation and purification (Scale bar: 5 µm). (D) Representative cryo-transmission electron microscopy (cryo-TEM) image of a single isolated biomimetic EV, showing an intact, unilamellar structure that indicates structural integrity post-isolation (Scale bar: 50 nm). (E) Size profile analysis of isolated vesicles by dynamic light scattering at different UV irradiation durations (3, 10, and 20 minutes), demonstrating a consistent size distribution (average diameters: 63±22 nm, 63±18 nm, and 65±22 nm, respectively) and confirming that the photocleavable linkers do not alter EV size or stability. (F) Schematic of a lipid fusion assay designed to assess the functionality of v-SNARE proteins in isolated donor (V) vesicles containing v-SNARE proteins and FRET-quenching fluorescent lipids RHOD-PE and NBD-PE. Subsequent mixing with target (T-SNARE) vesicles allows monitoring of v-SNARE functionality post-fusion, with restoration of NBD-PE fluorescence. (G) Quantification of membrane fusion efficiency using the FRET assay. Vesicles released by UV irradiation for 1 minute show fusion efficiency comparable to the ultracentrifugation technique, demonstrating that the isolated vesicles retain their biological activity. Error bars represent standard deviation (n=3).

For extrusion-based bEV fabrication, we mixed isolated EVs with freeze-thawed lipid vesicles composed of DOPC doped with 0.5% NBD-PE fluorescent lipid, and then extruded the mixture through polycarbonate membranes to generate discrete size populations. Size characterization by dynamic light scattering (DLS) and nanoparticle tracking analysis (NTA) showed consistent and reproducible control over vesicle diameter. DLS data showed mean diameters of 84 ± 12 nm and 146 ± 22 nm for vesicles extruded through 50 nm and 200 nm membranes, respectively, while NTA measured corresponding means of 96 nm and 149 nm (Fig. SI.1). When these two populations were combined to create a multimodal input sample, DLS and NTA again agreed closely (DLS: 136 ± 25 nm; NTA: 130 nm), confirming consistency across orthogonal sizing modalities (Fig. SI.1).

To simulate large-volume isolation conditions relevant to scalable workflows, the bEV populations were diluted into 50 mL buffer and processed using the platform. After 15 min capture, the beads were collected and the vesicles were released by UV irradiation (90 s) to induce photocleavage, eliminating the need for competitive D-biotin elution. DLS analysis of the photoreleased vesicles showed size distributions centered around 100 nm and 130 nm (Figure 2B), indicating that vesicle morphology was preserved throughout capture and release. Consistent with these measurements, single-vesicle fluorescence imaging revealed bright, diffraction-limited puncta (Figure 2C), and cryo-EM showed intact, spherical vesicles with well-defined bilayers (Figure 2D). As an additional integrity check, we also performed an experiment with detergent, which abolished vesicle fluorescence and confirmed that the recovered particles were membrane-enclosed vesicles rather than non-vesicular aggregates.To verify whether our probes promote vesicle clustering and aggregation, we incubated the bEV samples extruded through 50 nm membranes with probes for varying durations (3, 10, and 20 minutes) and analyzed by DLS. Across all incubation times, size distributions remained stable (average diameters 63 ± 22 nm, 63 ± 18 nm, and 65 ± 22 nm, respectively; Figure 2E), demonstrating that PLN exposure does not measurably increase vesicle size or polydispersity. These results support the physical robustness of the PLN design for vesicle processing and purification.

We then tested whether photorelease preserves membrane protein functionality using a v-SNARE–driven lipid-mixing fusion assay (Figure 2F). Donor bEVs (V) containing VAMP2 (v-SNARE) were labeled with NBD-PE/Rhodamine-PE to establish a FRET pair (fluorescence “OFF” state). These donor vesicles were prepared by detergent-assisted reconstitution: a dried lipid film was hydrated and mixed with isolated EVs in the presence of detergent and VAMP2, followed by overnight dialysis to remove detergent and yield proteo-bEVs with reconstituted SNAREs. Following PELA capture and UV-triggered release, donor vesicles were mixed with unlabeled acceptor vesicles (T) containing complementary t-SNAREs. Fusion-driven lipid mixing disrupts the FRET pair, restoring NBD fluorescence (“ON” state) as a quantitative readout of SNARE activity. Importantly, vesicles exposed to 1 min UV irradiation retained fusion efficiency comparable to density gradient–purified controls (Figure 2G), indicating that the isolation and photorelease conditions preserve functional membrane proteins. Collectively, these data demonstrate that the platform enables single-step engineering and isolation of bEVs without ultracentrifugation while preserving vesicle morphology, membrane integrity, and membrane protein function. The ability to recover structurally intact and biologically active vesicles at scale provides a strong foundation for downstream applications in nanomedicine, including drug delivery, gene therapy, and diagnostic profiling, where both purity and functional integrity are essential.

### 2.3 Orientation-Specific Enrichment of Nano Plasma Membrane Vesicles

Cell-derived plasma membrane nanovesicles are commonly used as biomimetic drug delivery systems, yet standard preparation methods are inefficient because they produce vesicles with mixed membrane orientations.[23, 24] Right-side-out (RSO) nanovesicles preserve native topology, presenting extracellular glycans and receptors on the exterior surface, whereas inside-out (ISO) vesicles invert this topology and expose cytosolic domains outward. Because correct protein orientation is essential for receptor-mediated targeting, immune evasion, and biological recognition, highly RSO-enriched nanovesicle preparations are a prerequisite for reliable drug delivery and biomimetic function. Conventional extrusion-based methods, however, produce vesicles with mixed orientations and provide no mechanism for selective enrichment.

To quantitatively assess and control membrane orientation, we selected red blood cell (RBC) membranes as a clinically relevant and scalable model system. RBC-derived vesicles are attractive delivery carriers due to their biocompatibility, immune evasion properties, and long circulation lifetimes. As illustrated in Figure 3A, micron-scale plasma membrane vesicles (RBC ghosts) were generated by hypotonic lysis and subsequently extruded through 100 nm polycarbonate membranes to form nano plasma membrane vesicles (nPMVs). Confocal microscopy of the micron-scale RBC ghosts labeled with NBD-PE revealed uniform membrane staining and giant unilamellar vesicle-like morphology (Figure 3B). We performed size analysis using the line profile (Figure 3C), yielding an average diameter of 3.87 µm (Figure 3D). We then quantified membrane orientation using a dithionite quenching assay, which selectively quenches NBD-PE exposed on the outer leaflet due to the membrane-impermeability of dithionite. In the unextruded micron-scale vesicles, nearly 100% fluorescence quenching was observed (Figure 3E), confirming preservation of native plasma membrane orientation. In contrast, after extrusion to the nanoscale, only ∼57% of nPMVs retained RSO orientation, with ∼43% adopting an ISO configuration (Figure 3F). These results confirm that extrusion-driven membrane fragmentation and resealing produce a statistically mixed-orientation population, underscoring the need for an active enrichment strategy.

**Figure 3.**
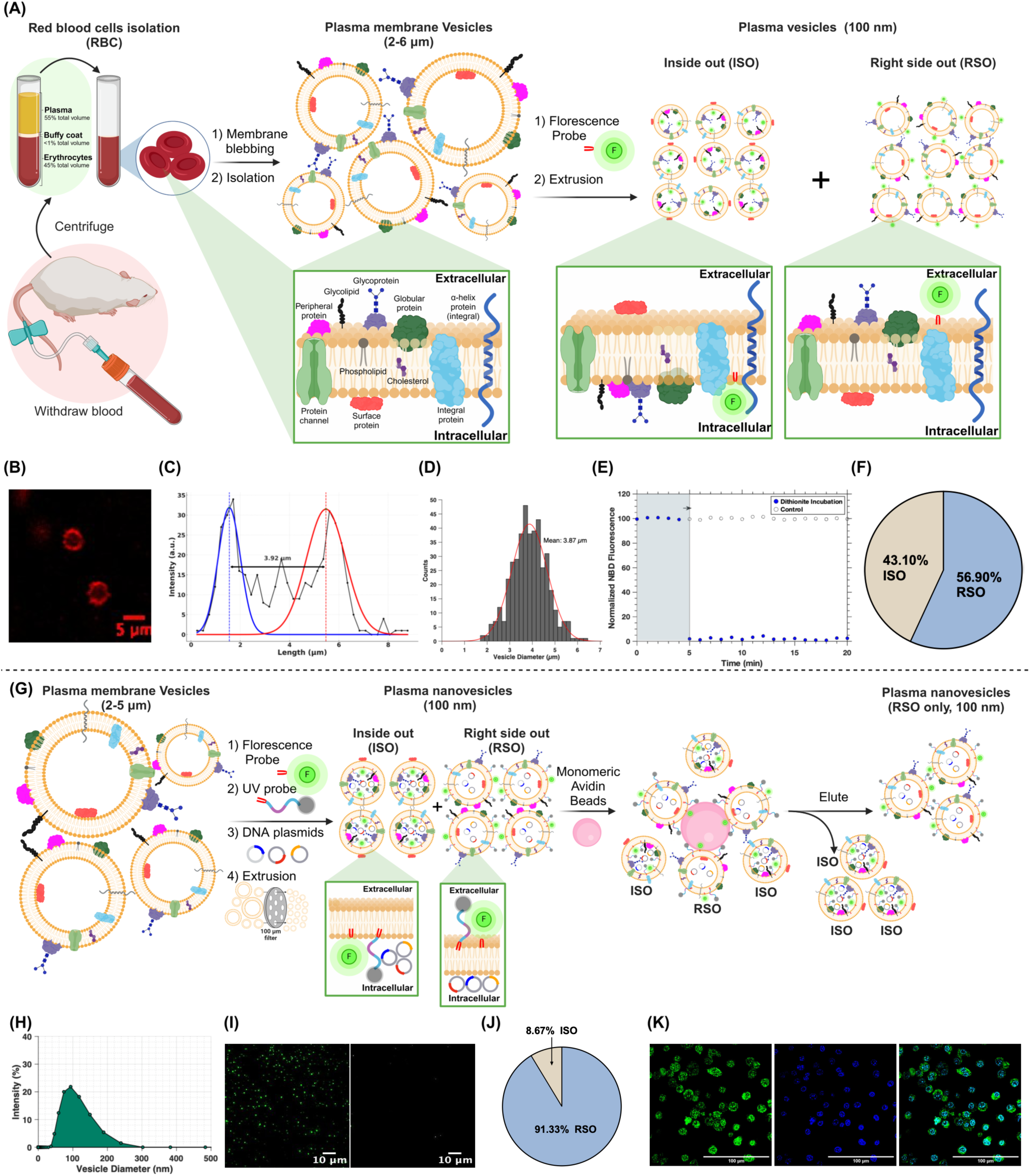
Isolation and Characterization of Red Blood Cell (RBC)-Derived Plasma Membrane nanoVesicles (PMnVs) with Defined Orientations and Sizes Using PLN Probes. (A) Schematic overview of the generation of PMnVs with inside-out (ISO) and right-side-out (RSO) orientations. Microvesicles (2-6 µm) are generated from isolated RBCs via membrane blebbing. Nanovesicles are produced through sequential extrusion of the microvesicles. (B) Representative fluorescence microscopy image of RBC membrane-derived microvesicles labeled with an NBD-PE membrane dye. Length (C) and size (D) profile analysis of isolated microvesicles. Dithionite incubation destroys the NBD-PE fluorescence of the outer leaflet only. Sodium dithionite at a concentration of 20 mM (solid circles) or buffer (open circles) was added after 5 min at 37 °C to a solution of vesicles containing NBD-PE. The complete loss of the signal after the addition of dithionite shows that all the fluorescence is present in the outer leaflet (E) but after extrusion the ratio becomes 43% and 57% indicating ISO and RSO distribution (F). (G) Schematic of the strategy for purifying RSO nanovesicles as a drug delivery vector using the PLN probe. (H) DLS analysis of purified RSO nanovesicles, showing a size distribution of around 100 nm. (I) Fluorescence microscopy images of PMnV containing NBD-PE fluorescent membrane dye (Scale bar: 5 µm). (J) Quantification of purified RSO nanovesicles using a dithionite quenching assay, revealing an enrichment of RSO vesicles to 83%. (K) Fluorescence microscopy images of purified RSO nanovesicles carrying VAMP2-GFP protein expressed in macrophage cell culture demonstrating the RSO orientation, retention of cargo, and potential for delivery to cells (Scale bar: 20 µm).

To selectively isolate RSO nanovesicles, we employed a leaflet-specific probe insertion strategy. By inserting functionalized lipid probes into the outer leaflet of the plasma membrane prior to extrusion, only vesicles that retain RSO topology after extrusion display the affinity tag on their external surface. In ISO vesicles, the tag becomes sequestered within the vesicle lumen and is inaccessible. After extrusion of RBC ghosts, nPMVs were immediately incubated with monomeric avidin–functionalized microbeads to capture vesicles presenting externally accessible affinity tags (Figure 3G). The moderate binding affinity of monomeric avidin enables stringent washing, effectively removing ISO vesicles while retaining RSO vesicles. Bound vesicles were subsequently eluted using excess free biotin to yield an RSO-enriched fraction. The isolated RSO vesicles exhibited a size distribution centered at ∼100 nm, as measured by DLS (Figure 3H). Fluorescence microscopy revealed bright, punctate vesicles that remained susceptible to dithionite quenching (Figure 3I), confirming the presence of an accessible outer leaflet. Quantitative dithionite analysis showed that 91% of the purified vesicles exhibited RSO orientation (Figure 3J), representing a substantial enrichment relative to the initial ∼57% RSO population. The small residual ISO fraction likely reflects minor probe insertion into inner leaflets due to multiple pass through the membranes during extrusion.

Finally, we evaluated whether orientation-specific isolation preserves functional cargo delivery capability. nPMVs loaded with VAMP2-GFP mRNA were subjected to the same RSO enrichment workflow and incubated with RAW264.7 macrophages. Confocal microscopy after 24 h revealed robust intracellular expression of VAMP2-GFP, confirming successful cytosolic delivery of functional RNA cargo (Figure 3K). This proof-of-concept demonstrates that RSO-enriched nanovesicles retain biological functionality and are competent for nucleic acid delivery. Collectively, these results establish a generalizable and scalable strategy for orientation-specific enrichment of plasma membrane nanovesicles.

### 2.4 Curvature-Encoded Photochemical Programming of Lipid Vesicle Populations

We next explored whether the PLNs could be used not merely as a capture–release element, but as a means to program vesicle behavior using light by exploiting intrinsic membrane properties. Lipid vesicles, including EVs and bEVs, exhibit continuous heterogeneity in size, curvature, and membrane organization. Rather than attempting to eliminate this heterogeneity through physical separation, we sought to determine whether it could be encoded into probe density and decoded through a controllable photochemical stimulus. Because amphiphilic PLNs integrate into lipid bilayers in a curvature-dependent manner, vesicles with different membrane curvatures intrinsically acquire different numbers of probes. This establishes a physical link between vesicle geometry and photochemical response. We hypothesized that controlled UV irradiation could therefore be used to differentially address vesicle subpopulations in a time-resolved manner, using light as an analog control parameter rather than a binary trigger.

PLNs were incorporated into EVs and bEVs using the workflow shown in Figure 4A. Vesicle populations were subsequently subjected to specified UV irradiation durations, and the vesicles released were examined using DLS. We observed that vesicle release occurred progressively as a function of irradiation time (Figure 4B), revealing a continuous photochemical response rather than an abrupt transition. Smaller diameters dominated vesicles released at early irradiation times, whereas prolonged exposure resulted in the release of larger vesicles (Figure 4C). Quantitative analysis of the DLS profiles using the area under the curve (AUC) demonstrated a amount of increase in vesicle release with irradiation time (Figure 4D), consistent with progressive photochemical cleavage of PLNs. This response arises from fundamental differences in probe density: highly curved membranes incorporate fewer probes and therefore require fewer cleavage events to disengage, whereas less curved membranes require longer exposure. We observed that this response is reproducible across different vesicle systems, indicating that it arises from fundamental membrane-probe interactions rather than vesicle-specific compositions.

**Figure 4.**
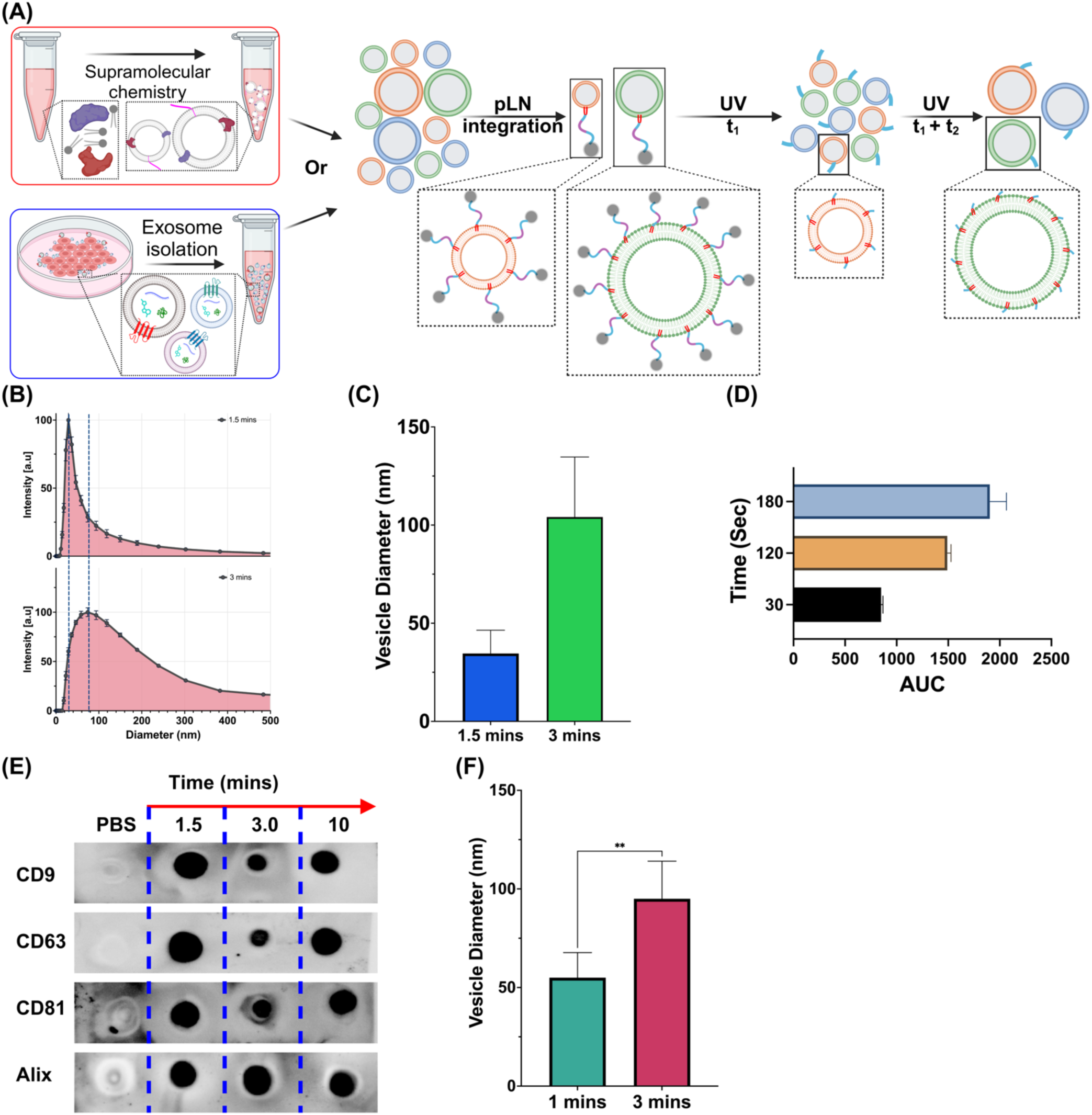
Light-Controlled Size-Based Separation of Vesicles Using Photocleavable Lipid Nanoprobes (PLNs). (A) Schematic illustration of PLN integration into the membranes of vesicles, either synthetically assembled (top) or isolated EVs (bottom). Subsequent UV irradiation triggers PLN cleavage, with the extent of cleavage dependent on light intensity, exposure time, and the number of PLNs per vesicle. This differential cleavage allows for size-based separation. (B) Representative Nanoparticle Tracking Analysis (NTA) profiles of vesicles after UV irradiation for 1.5 minutes (top) and 3 minutes (bottom), demonstrating a shift in the size distribution as a function of exposure time. (C) Quantification of vesicle size (diameter) from NTA data after 1.5 minutes and 3 minutes of UV irradiation. Smaller vesicles, with fewer integrated PLNs, undergo more rapid changes in size due to faster PLN cleavage. Error bars indicate standard deviation. (D) Bar graph comparing the area under the curve (AUC) of the NTA size distribution profiles after different UV irradiation times, representing the relative abundance of vesicles released. Changes in AUC reflect the light-induced separation. (E) Dot blot analysis of exosomal markers (CD9, CD63, CD81, and Alix) in vesicles after different durations of UV irradiation (1.5, 3, and 10 minutes in PBS), confirming the exosomal origin of the vesicles and showing that the markers are largely unaffected by the light exposure under these conditions. (F) Quantification of biomimetic EVs size after 1 minute and 3 minutes of UV exposure, demonstrating that smaller vesicles are released in response to shorter UV exposure due to their lower PLN density, while larger vesicles require longer exposure for comparable changes. Statistical significance was determined using a t-test.

Next, we tested photochemical programming of vesicle release does not compromise vesicle integrity or molecular identity. Dot blot analysis of canonical exosomal markers (CD9, CD63, CD81, and Alix) showed consistent expression across all irradiation conditions (Figure 4E), demonstrating that PLN cleavage and light exposure preserve vesicle protein content. Statistical analysis of vesicle diameters after short (1 min) and extended (3 min) irradiation further confirmed a significant and controllable shift in vesicle populations as a function of light dose (Figure 4F).

Beyond demonstrating photochemical control, this behavior has important implications for addressing EV heterogeneity. EV populations are intrinsically diverse, and conventional isolation methods often average over this diversity or impose rigid cutoffs based on size or density. In contrast, the PLN platform leverages membrane curvature as an intrinsic materials parameter, transducing it into probe density and enabling time-resolved access to vesicle subpopulations using a single external stimulus. Rather than eliminating heterogeneity, this strategy provides a programmable means to interrogate and access it while preserving vesicle integrity and biological functionality. Collectively, these results establish a generalizable framework for curvature-encoded photochemical programming of lipid vesicles. By transforming light into a continuous control parameter, this approach enables dynamic, non-destructive addressing of heterogeneous vesicle populations..

### 2.5 Isolation of High-Purity EVs from Human Plasma Samples for Clinical Applications

Having established photochemical control over vesicle capture, integrity, orientation, and heterogeneity, we next evaluated the translational potential of the platform using human plasma samples. Plasma is among the most challenging biological matrices for EV isolation due to its high abundance of lipoproteins, protein aggregates, and soluble components that overlap in size and density with EVs. Efficient recovery of high-purity, antigen-intact EVs from plasma remains a major bottleneck for clinical diagnostics and biomarker discovery.

We collected fresh human plasma samples from cognitively normal individuals and patients with Alzheimer’s disease in accordance with standard ethical protocols (Figure 5A). These preprocessed plasma samples were then incubated directly with PLNs, thereby facilitating spontaneous probe insertion into vesicle membranes. After thorough washing to eliminate unbound plasma components, vesicles were released through light-triggered photocleavage, which avoids harsh elution methods or competitive displacement strategies. This gentle isolation process is particularly important for engineering EVs intended for personalized medicine, where preservation of membrane integrity and surface functionality is essential.

**Figure 5.**
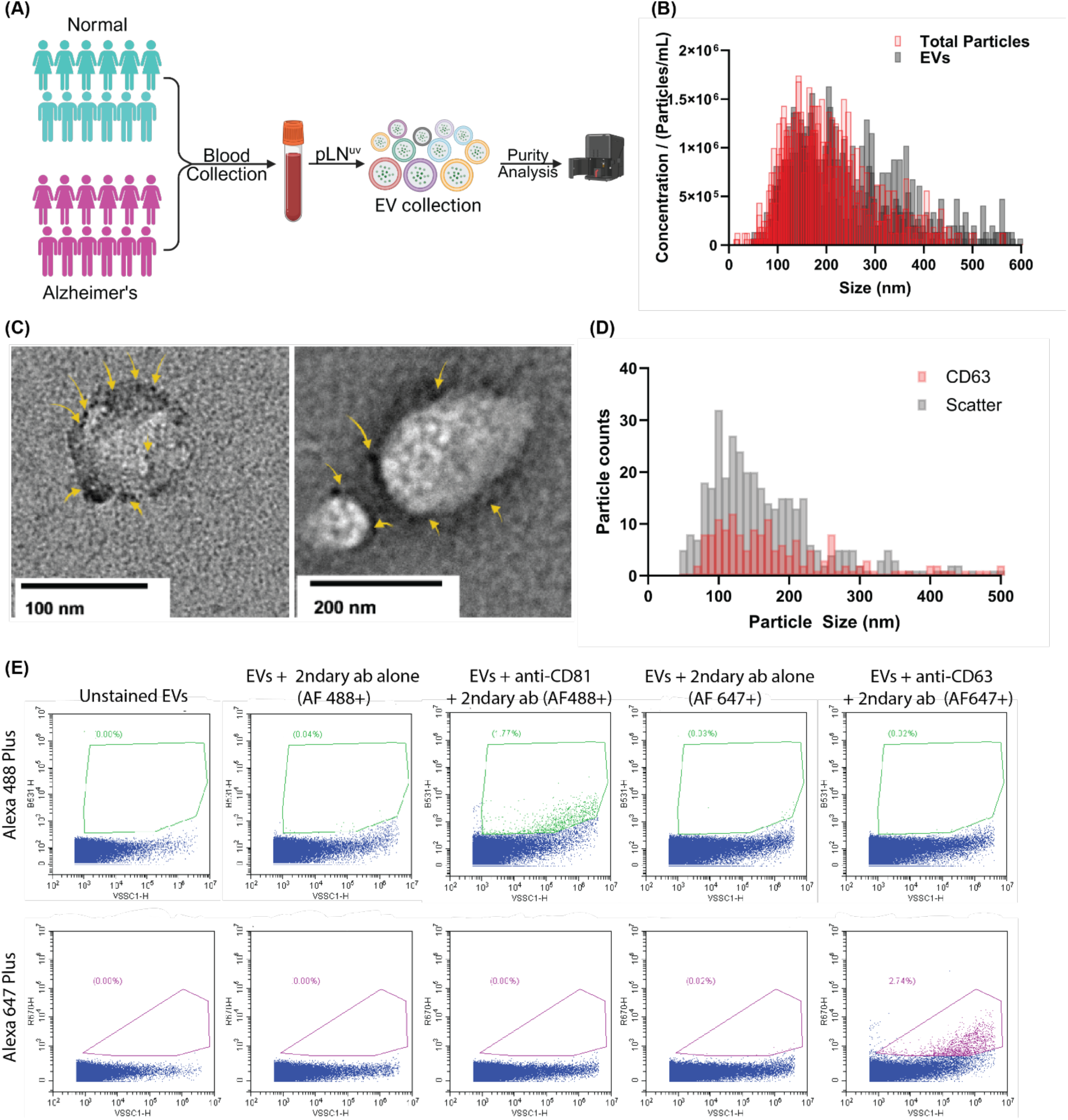
Rapid isolation and molecular profiling of plasma EVs. (A) Schematic of clinical workflow illustrating donor recruitment, blood processing, and PELA-based EV purification. (B) NTA profiles showing depletion of non-vesicular particles and enrichment of sEV-sized vesicles. (C) TEM images of isolated plasma EVs labeled with CD63 antibody containing gold nanoparticles, displaying intact unilamellar morphology with minimal background contamination. (D) Nanoscale flow cytometry demonstrating CD63⁺ EV subpopulation relative to total scatter events. (E) Representative scatter plots of EV immunolabeling using secondary antibodies, confirming preservation of vesicle surface markers and low nonspecific background.

To independently verify vesicle purity, isolated particles were labeled with MemGlow, a fluorogenic membrane-intercalating dye that selectively reports lipid bilayers. Because MemGlow does not label protein and self-quenches in solution, fluorescence directly reflects membrane-enclosed vesicles. Quantitative analysis showed that >95% of recovered particles were MemGlow-positive, confirming high EV purity. The NTA analysis showed a reduction in non-vesicular particles after photorelease, with a clear enrichment of vesicles in the 80-180 nm size range characteristic of small EVs (Figure 5B). Transmission electron microscopy (TEM) further confirmed the presence of intact, unilamellar vesicles that were immunogold-labeled for CD63, demonstrating that the recovered vesicles retained accessible surface markers and preserved membrane topology (Figure 5C). To rigorously validate vesicle identity and antigen preservation, we performed nanoscale flow cytometry targeting CD63, a canonical EV surface marker. Particle-by-particle analysis revealed a distinct CD63⁺ vesicle subpopulation within the expected size window, clearly separated from scatter-only events (Figure 5D). We further assessed the compatibility of the isolated vesicles with antibody-based labeling workflows relevant to clinical diagnostics. Comprehensive immunophenotyping using fluorescently labeled antibodies showed low nonspecific background and clear discrimination between labeled and unlabeled vesicles across conditions (Figure 5E). These results indicate that vesicle surface proteins remain structurally intact and functionally accessible following light-triggered release, enabling reliable antibody binding and detection. Collectively, these findings demonstrate that the platform enables rapid, high-purity isolation of antigen-intact EVs directly from complex human plasma samples. The ability to recover clinically relevant EVs with preserved membrane topology and immunoreactivity positions this platform as a powerful materials-enabled solution for liquid biopsy applications, diagnostic profiling, and translational EV research.

Together, these data establish that the PLD platform enables rapid, high-purity isolation of EVs directly from human plasma, preserving vesicle morphology, molecular markers, and analytical compatibility. This capability underscores the translational potential of PLNs for clinical-grade vesicle preparation and blood-based biomarker discovery in neurodegenerative disease and other disorders.

## 3. Conclusion

This work establishes a programmable, light-responsive materials platform that redefines how lipid nanovesicles can be isolated, fractionated, and functionally preserved at scale. By integrating PLN with reversible electrostatic capture, we transform vesicle purification from a passive separation process into an active, materials-encoded operation in which membrane curvature, orientation, and probe density are directly coupled to a controllable photochemical stimulus. This design enables rapid (<15 min), high-purity isolation of extracellular vesicles and plasma membrane nanovesicles while preserving native membrane architecture, protein functionality, and cargo integrity.

Beyond rapidity and scalability, the platform introduces two capabilities that are largely inaccessible with existing technologies: orientation-specific enrichment of right-side-out plasma membrane vesicles and curvature-encoded, light-programmed access to heterogeneous vesicle subpopulations. In this framework, light functions not as a binary release trigger, but as a continuous analog control parameter that decodes intrinsic membrane properties into selective vesicle release. This approach allows heterogeneity to be harnessed as a functional design variable. Importantly, the platform operates across vesicle classes, from biomimetic assemblies to native EVs isolated directly from human plasma, and remains compatible with downstream biochemical, imaging, and immunophenotyping workflows. The ability to rapidly generate antigen-intact, orientation-defined, and size-resolved vesicle populations addresses a central bottleneck in vesicle-based diagnostics, drug delivery, and standardized manufacturing.

More broadly, this research treats photocleavable lipid nanoprobes as functional soft-matter components capable of encoding geometry, topology, and release dynamics within biological membranes. Combining membrane physics with photochemistry, the platform offers a versatile approach for actively controlling lipid assemblies through external stimuli. This not only provides a practical method for clinical-grade vesicle separation but also serves as a conceptual framework for creating responsive, information-encoded nanomaterials at biointerfaces.

## 4. Materials & Methods

### 4.1 Materials and Reagents

Synthetic phospholipids, including 1,2-dioleoyl-sn-glycero-3-phosphocholine (DOPC), 1,2-dioleoyl-sn-glycero-3-phosphoethanolamine (DOPE), 1,2-dioleoyl-sn-glycero-3-phospho-L-serine (DOPS), and cholesterol, were purchased from Avanti Polar Lipids. Fluorescent lipid analogs N-(7-nitrobenz-2-oxa-13-diazol-4-yl)-1,2-dipalmitoyl-sn-glycero-3-phosphoethanolamine (NBD-PE) and Lissamine-Rhodamine-PE (Rh-PE) were obtained from Avanti Polar Lipids. Biotinylated Poly-L-lysine (20 units) was obtained from Nanosoft Polymers. D-biotin, sodium dithionite, and Triton X-100 were purchased from Sigma-Aldrich. Monomeric avidin agarose beads (High Capacity) were purchased from Thermo Scientific. The fluorogenic membrane dye MemGlow-488 was obtained from Cytoskeleton, Inc. Recombinant VAMP2 (v-SNARE) and Syntaxin-1A/SNAP-25 (t-SNARE) proteins were expressed and purified from E. coli as described previously.[25, 26] Non-photoreactive chemical probes were purchased from BroadPharm and Nanocs. All other reagents were analytical grade and used as received.

### 4.2. Synthesis of Photocleavable Lipid Nanoprobe (PLN)

The PLN construct was synthesized by conjugating a hydrophobic DSPE lipid anchor to a biotin-PEG3 affinity tag. Briefly, 1,2-distearoyl-sn-glycero-3-phosphoethanolamine (DSPE, 1 equivalent) and photocleavable biotin-PEG3-NHS ester (1 equivalent) were dissolved in anhydrous dichloromethane (4 mL) in the presence of triethylamine (TEA, 1.5 equivalent) in an oven-dried round bottom flask (25 mL). The mixture was stirred at room temperature for 24 hours and monitored with thin-layer chromatography (TLC). The solvent was then removed under reduced pressure. The crude residue was purified by automated normal phase chromatography (dichloromethane/methanol 20/80) to obtain a white powder.

### 4.3. Cell Culture

Rat insulinoma INS-1 cells were cultured in a humidified incubator at 37 °C with 5% CO₂ in complete RPMI-1640 medium (containing 11 mM glucose) supplemented with 10% fetal bovine serum (FBS), 2 mM L-glutamine, 10 mM HEPES, 1 mM sodium pyruvate, 50 μM β-mercaptoethanol, and 1% penicillin/streptomycin. They were grown on standard tissue culture-treated plastic without additional coating and passaged at approximately 70-80% confluence. To collect conditioned medium for EV isolation, cells were rinsed with pre-warmed PBS and incubated for 24 hours in the same medium, prepared with EV-depleted FBS. The conditioned medium was then collected and filtered through a 0.22 µm PES low protein-binding filter to remove debris before proceeding with EV capture.

### 4.4. EV Isolation

EV isolation was performed using High-Capacity Avidin-Biotin (HCAB) affinity chromatography following established protocols. Avidin-HCAB slurry (200 μL, Thermo Scientific) was loaded into 0.8 mL centrifuge columns equipped with 30 μm polyethylene filters (Thermo Scientific), with larger volumes processed using 5-10 mL columns. The 4% beaded agarose resin (50% slurry) was buffer-exchanged through five 200 μL washes with Buffer A. EVs were treated with 100 μg LN in 50 mL culture medium. Buffer-exchanged HCAB was combined with LN-labeled samples and agitated for 30 minutes or overnight for enhanced capture. Captured vesicles were washed 5-10 times with buffer A. Elution was achieved using either: (1) competitive displacement with 100 μL of 2.5 mM D-biotin for 2 minutes, or (2) photocleavage by exposing columns to 365 nm UV (5 mW/cm²) without biotin. Eluted vesicles were recovered by centrifugation (200 g, 20 seconds). HCAB resin was regenerated using 0.1 M glycine (pH 2.8) followed by buffer equilibration for reuse.

### 4.5. Biomimetic Extracellular Vesicle (bEV) Fabrication

Lipid mixtures containing DOPC: DOPE: DOPS: Cholesterol (39:20:10:20 mol%) with 1% NBD-PE (for imaging) were mixed in a borosilicate glass culture tube and dried under a nitrogen stream for 15 minutes. Traces of solvent were removed by vacuum desiccation for 1 hour. The individual lipid stock solutions are purchased from Avanti as chloroform stocks. The dried lipid films were rehydrated with 1 mL of 25 mM HEPES, 150 mM KCl, pH 7.4 to a final lipid concentration of 3 mM. Buffers were filtered using a syringe 0.22 µm size syringe filter and equilibrated to room temperature prior to use. The resulting vesicles were combined with isolated native EVs and extruded through 50 nm or 200 nm pore size polycarbonate membranes to generate defined bEV size populations. During preparation, samples and glassware were protected from ambient light (foil-wrapped) to minimize photodegradation of fluorescent and photocleavable components.

### 4.6. Constructing bEVs with Transmembrane Proteins

V-SNARE (VAMP2) and t-SNARE proteins were purified as described. Isolated native EVs from cell culture were resuspended in Buffer A and mixed with v-SNARE proteins and dried lipid layer NBD-PE and Rhod-PE lipids with 1% (w/v) octyl-β-D-glucoside (OG). After mixing them for 40 minutes, the mixture was diluted 2 times with non-OG buffer to achieve rapid dilution. The mixture was dialyzed overnight with Bio-Beads SM-2 (Bio-Rad). Post-dialysis, samples were mixed with 150 μL of 80% OptiPrep (iodixanol), loaded into 0.8-mL ultracentrifuge tubes (Beckman Coulter), and overlaid with 250 μL of 30% OptiPrep and 50 μL Buffer A. Gradient centrifugation was performed at 48,000 rpm for 4 hours using an SW55 rotor. Proteo-bEVs were collected between the buffer and 30% OptiPrep interface.

### 4.7. Lipid Mixing Assay

Membrane fusion was monitored using a FRET-based lipid mixing assay adapted from established protocols. V-SNARE proteo-bEVs containing NBD-DOPE (donor) and Rhodamine-DOPE (acceptor) were mixed with unlabelled t-SNARE proteoliposomes. Both populations were maintained at a total lipid concentration of 3 mM and preincubated at 37°C before being combined in 96-well white polystyrene plates. NBD fluorescence was measured at 460/535 nm (excitation/emission) at 1-minute intervals for 100 minutes using a microplate reader. Data were normalized as: % Lipid Mixing = 100% × (Ft - F0)/(Fmax - F0), where Ft is fluorescence at time t, F0 is initial fluorescence, and Fmax represents maximum fluorescence following complete lipid mixing with 2.5% (w/v) n-dodecylmaltoside.

### 4.8. Isolation of Red Blood Cells from Mice

Mouse blood samples were collected via tail vein puncture following Yale IACUC-approved protocols. Mice were briefly warmed under a heat lamp or in warm water (37°C) to dilate tail veins. The tail was disinfected with 70% ethanol and a small incision was made at the lateral tail vein using a sterile scalpel blade or 25-gauge needle. Blood (200-500 μL) was collected directly into tubes containing 10 μL of 0.5 M EDTA per mL of anticipated blood volume to prevent coagulation. Hemostasis was achieved by applying gentle pressure with sterile gauze. Samples were processed within 30 minutes of collection while maintained on ice. Whole blood was centrifuged at 500 × g for 10 minutes at 4°C using a refrigerated centrifuge. The plasma supernatant and buffy coat layers were carefully removed by aspiration, retaining the red blood cell pellet. Red blood cells underwent three sequential washes to remove plasma proteins and cellular debris. Each wash consisted of gentle resuspension in 10× volume of sterile PBS followed by centrifugation at 500 × g for 10 minutes at 4°C. After the final wash, cells were resuspended in minimal volume PBS and assessed for purity and viability. This yielded approximately 100-200 μL packed RBCs from 0.5 mL whole blood for immediate downstream processing.

### 4.9. Preparation of RBC Ghosts and Inside-Out RBC Membrane Vesicles (IOVs)

Freshly isolated mouse RBCs were washed three times in cold PBS (500 × g, 10 min, 4 °C) to remove plasma proteins. The derivation of RBC-membrane vesicles follows a previously reported methods with slight modifications.[24, 27] Briefly, RBC ghosts were generated by hypotonic lysis to remove cytosolic hemoglobin while preserving membrane structure. Washed RBCs were diluted 1:20 (v/v) into ice-cold hypotonic lysis buffer (5 mM sodium phosphate, pH 8.0, 1 mM EDTA) and incubated on ice for 30 min with gentle inversion. Membranes were pelleted (20,000 × g, 15 min, 4 °C), and the supernatant was discarded. The pellet was resuspended in fresh hypotonic buffer and the wash/lysis cycle was repeated for 5 times until the membrane pellet appeared pale and the supernatant was nearly colorless. Final ghosts were resuspended in Buffer A. RBC ghosts were labeled by adding 18:1 NBD-PE (6 µL of a 1 mg/mL stock; 6 µg, ∼6.5 nmol) as a dried lipid film, followed by resuspension with the ghost suspension prior to extrusion. Based on erythrocyte phospholipid content and typical mouse RBC counts, this corresponds to approximately ∼0.5 mol% relative to total RBC membrane phospholipid. Labeled ghosts were extruded through a 100 nm polycarbonate membrane to generate RBC membrane vesicles. For enrichment of RSO vesicles, ghosts were incubated with the indicated PLN before extrusion. Following extrusion, vesicles were incubated with monomeric avidin agarose beads to capture probe-accessible vesicles. The unbound fraction was collected as the probe-inaccessible population, and bead-bound vesicles were washed thoroughly prior to downstream analysis and eluted where indicated using D-biotin. Samples were protected from light during all steps involving fluorescent reporters.

### 4.10. Dithionite Quenching Assay for Membrane Orientation

Leaflet accessibility was quantified by sodium dithionite quenching of NBD fluorescence. Vesicles were diluted in Buffer A and baseline fluorescence was recorded (Ex/Em 460/535 nm). Freshly prepared Sodium dithionite solution was added to a final concentration of 20 mM, and fluorescence was monitored to a stable plateau. n-Dodecyl β-D-maltoside was then added (final 0.1-0.5% v/v) to rupture membranes and define the residual background.

### 4.11. Isolation of EVs from Human Plasma

EV isolation from human plasma was performed using a modified affinity capture protocol. Lipophilic nano-biotin (LN, 0.5 mg) was dissolved in 100 μL Buffer A and combined with 100 μL avidin-HCAB slurry (Thermo Scientific) in a 1.5 mL microcentrifuge tube. The mixture was incubated at room temperature with agitation for 1 hour. Human plasma (250 μL) was subjected to sequential centrifugation: first at 2,000g for 20 minutes, then the supernatant was re-centrifuged at 10,000g for 20 minutes at room temperature to remove cellular debris and large particles. The clarified plasma supernatant was added to the PLN-HCAB mixture and incubated overnight at 4°C with rotation. Following overnight capture, the plasma-PLN-HCAB mixture was agitated for 30 minutes before transfer to 0.8 mL spin columns equipped with 30 μm filters. The columns were centrifuged at 1,000g for 30 seconds at 4°C, and flow-through was discarded. Captured EVs were washed ten times with 500 μL sterile Buffer A using identical centrifugation conditions. EV elution was achieved by applying 150 μL of 2.5 mM D-biotin to the washed resin, followed by 15-minute incubation with agitation. EVs were recovered by centrifugation at 1,500g for 1.5 minutes at 4°C into fresh collection tubes.

### 4.12. Labeling of Human Plasma EVs

For on-column staining, captured plasma EVs were distributed into filter tubes as separate aliquots for different labeling conditions. Primary antibodies anti-CD63(Proteintech), anti-CD81 (Proteintech) were incubated with captured EVs for 1 h at room temperature under agitation. After washing, fluorescent secondary antibodies were applied (typical dilution 1:50) for 1 h. For immunogold TEM workflows, a small aliquot was incubated overnight at 4 °C with a gold-conjugated secondary antibody. For membrane integrity readouts, MemGlow dye was incubated with the eluted EVs prior to downstream analysis.

### 4.13. Vesicle Size and Concentration Measurements

*<u>Dynamic Light Scattering (DLS):</u>* Hydrodynamic diameters were measured on a Wyatt DynaPro Nanostar. Eluted samples were loaded into disposable cuvettes and measured at 25 °C, typically as ten sequential acquisitions with short intervals between runs. Data were processed using regularization-based batch analysis to resolve size distributions and assess aggregation.

*<u>Nanoparticle Tracking Analysis (NTA):</u>* Size distributions and particle concentrations were measured on a ZetaView x30 (Particle Metrix) using a 405 nm laser. Instrument settings were kept constant across comparable runs. Samples were diluted to the optimal particle-per-frame regime and analyzed using ZetaView software (v8.04.04). Calibration was performed using polystyrene reference beads under manufacturer-recommended dilution.

### Single Vesicle Fluorescence Microscopy

Single vesicle imaging was performed using a custom total internal reflection fluorescence (TIRF) microscopy system based on previously described configurations.[28, 29] The system comprised a Nikon inverted microscope equipped with three laser excitation lines (488, 532, and 633 nm), a Photometrics DV2 dual-view system, and an Andor EMCCD camera for simultaneous dual-channel acquisition. All images were acquired at 512 × 512 pixel resolution with 16-bit dynamic range and standardized 150 ms exposure times across both detection channels. Image processing involved conversion to TIFF format using FIJI software, [30]followed by analysis using custom algorithms developed in Wolfram Mathematica. The analytical pipeline enabled automated vesicle identification and quantitative colocalization analysis. Colocalization efficiencies were calculated based on vesicle populations identified within each fluorescent channel, providing quantitative measures of multi-marker expression on individual vesicles.

### Negative Stain Electron Microscopy

Negative stain electron microscopy was performed to visualize gold-labeled sEV morphology following established protocols for extracellular vesicle imaging. CD63-labeled sEV samples were prepared by applying 10 μL of isolated vesicles directly onto glow-discharged carbon Type-B grids (400 mesh, copper; Ted Pella) and incubated for 15 minutes at room temperature. Sample preparation involved a two-step staining procedure: grids were initially rinsed with 5 μL of 2% (w/v) uranyl formate solution, followed by negative staining with an additional 5 μL of 2% uranyl formate for 1 minute. Excess stain was removed by blotting with filter paper. Imaging was performed using a JEOL JEM-1400Plus transmission electron microscope operating at 80 kV, equipped with a bottom-mounted 4k × 3k charge-coupled device camera (Advanced Microscopy Technologies). Images were acquired at appropriate magnifications to visualize individual vesicle morphology and gold-labeling distribution. This approach enabled assessment of vesicle integrity, size heterogeneity, and specific protein labeling efficiency at the single-particle level.

### Western Blot Analysis

EV protein characterization was performed using standard western blotting protocols adapted for extracellular vesicle analysis. EV samples were lysed in Laemmli sample buffer and denatured by boiling at 100°C for 10 minutes to ensure complete protein solubilization and denaturation. Protein separation was achieved by sodium dodecyl sulfate-polyacrylamide gel electrophoresis (SDS-PAGE) using appropriate acrylamide concentrations based on target protein molecular weights. Following electrophoretic separation, proteins were transferred to nitrocellulose or PVDF membranes according to Bio-Rad western blot protocols. Immunodetection was performed using primary antibodies specific to EV marker proteins. Blots were incubated with primary antibodies overnight at 4°C with gentle agitation, followed by appropriate secondary antibody incubation. Protein bands were visualized using enhanced chemiluminescence (ECL) western blot substrate or and imaged using standard chemiluminescent detection systems. This approach enabled quantitative assessment of EV-enriched proteins and confirmation of successful cellular component depletion.

### Statistics and Reproducibility

All data are presented as mean ± standard error of the mean (SEM) or mean ± standard deviation (SD) as specified in figure legends and text. Statistical analyses were performed using GraphPad Prism 10.0 software with appropriate tests selected based on experimental design and data distribution. For comparisons between two experimental groups, unpaired Student’s t-tests were employed. Multiple group comparisons were analyzed using two-way analysis of variance (ANOVA) followed by Tukey’s multiple comparisons test to control for family-wise error rates. Statistical significance was defined as P < 0.05 for all analyses. Significance levels are indicated using standardized notation: single asterisk (*) represents P ≤ 0.05; double asterisk (**) represents P ≤ 0.01; triple asterisk (***) represents P ≤ 0.001; quadruple asterisk (****) represents P ≤ 0.0001. All experiments were performed with appropriate biological and technical replicates as specified in individual figure legends, ensuring adequate statistical power for meaningful interpretation of results.

## Supporting Information

Supporting Information is available from the Wiley Online Library or from the author.

## Acknowledgements

We thank the WM Keck Biotechnology Resource Laboratory at the Yale School of Medicine for their assistance with the Zetaview NTA. We thank Particle Metrix for providing us with a demo of the ZetaView Evolution NTA.

